# High Throughput Mechanobiological Screens Enable Mechanical Priming of Pluripotency in Mouse Fibroblasts

**DOI:** 10.1101/480517

**Authors:** Jason Lee, Miguel Ochoa, Pablo Maceda, Eun Yoon, Lara Samarneh, Mitchell Wong, Aaron B. Baker

**Author notes:** Correspondence to: Aaron B. Baker, Ph.D., University of Texas at Austin, Department of Biomedical Engineering, 1 University Station, BME 5.202D, C0800, Austin, TX 78712.

## Abstract

Transgenic methods for direct reprogramming of somatic cells to induced pluripotent stem cells (iPSCs) are effective in cell culture systems but ultimately limit the utility of iPSCs due to concerns of mutagenesis and tumor formation. Recent studies have suggested that some transgenes can be eliminated by using small molecules as an alternative to transgenic methods of iPSC generation. We developed a high throughput platform for applying complex dynamic mechanical forces to cultured cells. Using this system, we screened for optimized conditions to stimulate the activation of Oct-4 and other transcription factors to prime the development of pluripotency in mouse fibroblasts. Using high throughput mechanobiological screening assays, we identified small molecules that can synergistically enhance the priming of pluripotency of mouse fibroblasts in combination with mechanical loading. Taken together, our findings demonstrate the ability of mechanical forces to induce reprograming factors and support that biophysical conditioning can act cooperatively with small molecules to priming the induction pluripotency in somatic cells.

---

Cellular reprograming is a process in which differentiated, mature cells are converted into induced pluripotent stem cells (iPSCs) through the expression of specific transcription factors. Generation of iPSCs is typically achieved through the expression of Oct-4, Klf-4, Sox2, and c-Myc (Takahashi and Yamanaka, 2006). The discovery of the means to reprogram mature cells to iPSCs and subsequently differentiate them into defined lineages has immense potential to revolutionize cell-based therapeutics, drug screening and scientific investigation of many diseases. However, current protocols for creating iPSCs are often inefficient and time consuming (Lujan et al., 2015; Polo et al., 2012). Moreover, genetic modification of cells to overexpress the transcription factors has the risk of creating mutations in the reprogrammed cells and tumorigenicity is a major concern with iPSC-based therapies (Ben-David and Benvenisty, 2011). Several groups have found that small molecules can replace some or all of the transcription factors that induce cellular reprograming (Hou et al., 2013; Ichida et al., 2009; Lin et al., 2009; Zhu et al., 2010), supporting that reprogramming is possible in the absence of genetic modification of the somatic cells.

Biophysical forces are now being recognized as important modulators of biological processes in many fields including cancer (Heldin et al., 2004; Jain et al., 2014), stem cell biology (Engler et al., 2006; Henderson et al., 2017; Lee et al., 2011; Yang et al., 2011) and embryological development (Goldie et al., 2008). Recent work has suggested a link between the mechanical environment and pluripotency (Ireland and Simmons, 2015). In addition, mechanical force can lead to chromatin remodeling and altered binding of transcription factors (Iyer et al., 2012; Tajik et al., 2016). Recent studies have also shown that application of mechanical strain can reduce the expression of pluripotency in mouse embryonic stem cells (Hazenbiller et al., 2017; Hazenbiller et al., 2018). In hPSCs, mechanical strain helped to maintain pluripotency through increased expression Nodal, TGF-β, and Activin (Saha et al., 2006, 2008). However, in other studies, mechanical strain reduced the expression of pluripotency transcription factors and signaling pathways related to pluripotency in iPSCs (Teramura et al., 2012). Mechanical stretch also enhanced the reprograming of cells treated with retrovirus-delivered pluripotency transcription factors without altering the efficiency of viral transduction (Kim et al., 2017). Other studies have linked alterations in pluripotency to changes in cell shape or substrate stiffness (Eroshenko et al., 2013; Macri-Pellizzeri et al., 2015; Maldonado et al., 2017). Thus, while the mechanical environment has a powerful effect on cellular reprogramming it is unclear how best to optimize the forces and/or environment to enhance pluripotency.

To investigate the role of applied mechanical forces in priming pluripotency in somatic cells, we developed a high throughput system for performing mechanobiological screens. This system allows the application of mechanical stretch to cells in a high throughput format of up to 576 wells individually, configured in removable multi-well plates that can be used in high content imaging systems and plate reading assays. Using this mechanobiological screening platform, we examined whether there were mechanical conditions that could prime somatic cells for developing pluripotency. Combining optimized mechanical conditioning with a drug screen of small molecule signaling modulators, we identified compounds that could further enhance the expression of pluripotency transcription factors in mouse fibroblasts. Our studies reveal that optimized mechanical stimulation combined with small molecule inhibitors can markedly enhance the pluripotency of somatic cells in the absence of transgene delivery.

## Results

### High throughput system for studying stem cell mechanobiology

We created a high throughput mechanical loading system that allows the application of mechanical stretch to cells cultured in six 96-well plates simultaneously. The system applies mechanical load by displacing an array of pistons mounted on a platen through a flexible bottom culture plate. The high throughput biaxial oscillatory strain system (HT-BOSS) drives the motion of a platen using a true linear motor (**Fig. 1A; Supplemental Fig. 1**). Teflon pistons are mounted on the platen that can be driven to displace a flexible culture surface within a custom culture plate (**Fig. 1B, C**). This system can apply strains based on the displacement of the piston and there is a linear relation of displacement to strain for a broad range of mechanical strains (**Fig. 1D**). The height of each piston can be adjusted individually, allowing calibration of each piston for accurate strain application (**Supplemental Fig. 2**). The system uses a true linear (voice coil) motor that allows customizable displacements to create complex dynamic strain waveforms. We verified that the system could apply the sine waveform and two physiologic strain waveforms derived from the displacement of the arterial wall in the aorta and brachial arteries (**Fig. 1E**) (Lee et al., 2013). The average shear stress on the culture surface scaled nearly linearly with frequency of loading and maximum strain (**Supplemental Fig. 3, 4**). The shear stress applied was very low and in the range of ~0.5 to 4 mPa over all of the conditions tested. In comparison to simulations of a larger format system with 35 mm diameter wells, the shear stress generated in this high throughput system were about 10-fold lower (6.35 mm diameter wells) (Lee and Baker, 2015; Lee et al., 2013).

**Figure 1.**
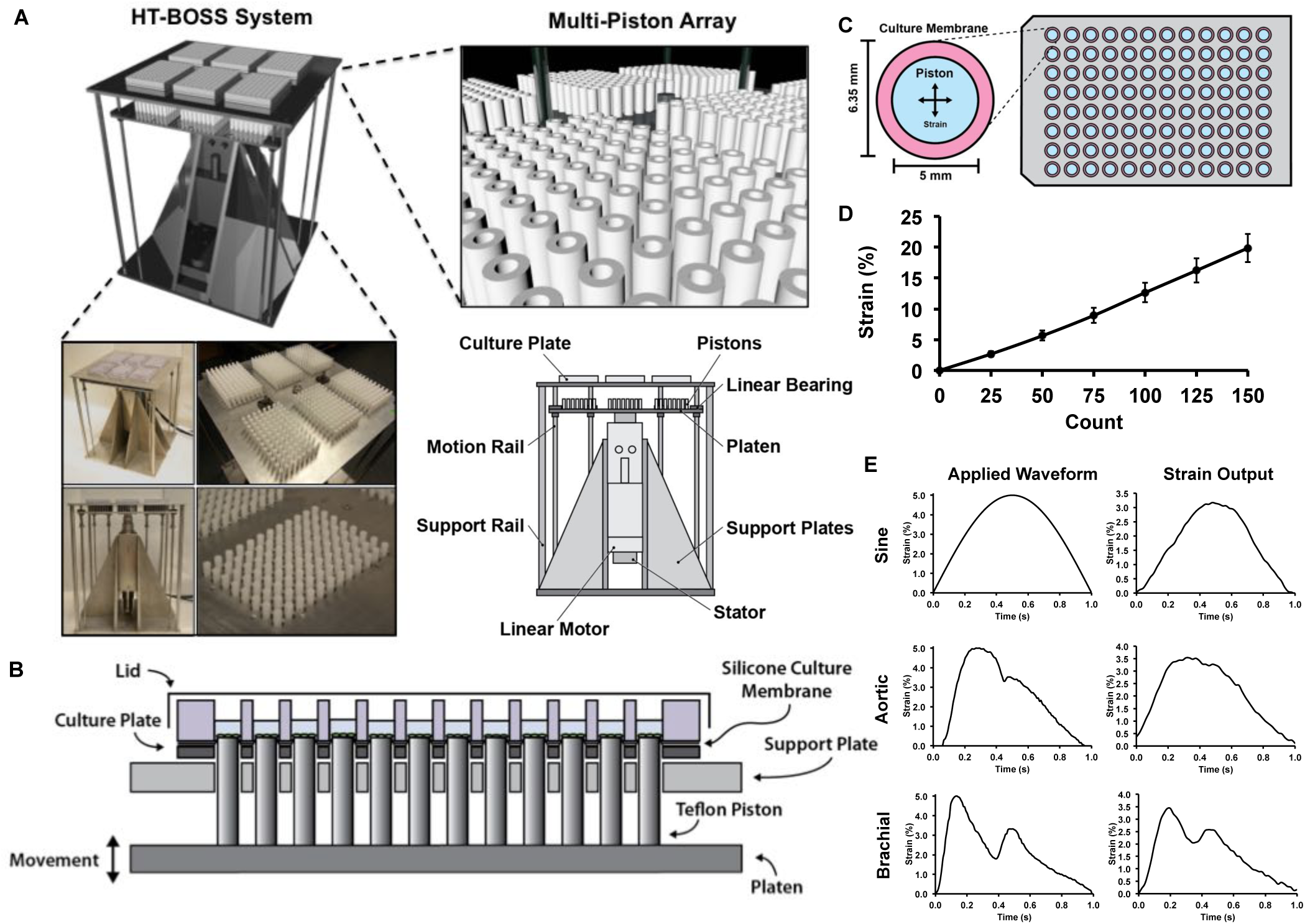
High throughput system for applying mechanical forces to cultured cells. (A) The system applies mechanical strain to cells cultured in a 96 well plate format. A platen with 576 pistons is moved by a linear motor to displace a flexible membrane in six 96-well plates with flexible culture surfaces. (B) The culture plate consists of two plates that connect to hold a thin silicone membrane that serves as a culture surface. Thicker silicone membranes serve to seal each well and prevent leakage between the wells. Teflon pistons mounted on the platen are displaced into the membrane to apply mechanical strain to the cells cultured in each well. (C) Top view of the system showing the relative geometry of the piston and the culture surface. (D) Average strain applied by the system with vertical displacement of the platen. Each motor count is 10 µm of vertical displacement. (E) Dynamic strain waveforms produced by the system through control of the platen motion with the linear motor. The aortic and brachial waveforms simulate the strain on the arterial wall during the cardiac cycle in the body.

### Multi-strain screening reveals that high level mechanical strains lead to increased expression of Oct-4, Sox2 and SSEA1

Using the high throughput loading system, we applied a range of load simultaneously by adjusting the heights of the pistons and calibrating the maximum strain applied (**Fig. 2A, B**). We applied mechanical strain to mouse embryonic fibroblasts (MEFs) that expressed an octamer-binding transcription factor 4 (Oct-4) eGFP transgene at 0.1 Hz with varying maximal strain from 0 to 17.5% strain. Using a plate reader, we assayed the expression of Oct-4 over seven days of loading (four hours of loading per day). We found that most levels of mechanical strain increased Oct-4 expression from approximately 1.5 to 2.5 fold (**Fig. 2C, D**). The highest levels of Oct-4 were observed in cells exposed to 17.5% strain. There was not a significant increase in Oct-4 expression over the days of loading (**Supplemental Fig. 5**). After 7 days of loading, we immunostained for pluripotency markers sex determining region Y-box 2 (Sox2) and Stage-specific embryonic antigen 1 (SSEA1). We found significant increases in all levels of mechanical strain for both markers and the greatest increase in markers for 17.5% mechanical strain (**Fig. 2E-G**).

**Figure 2.**
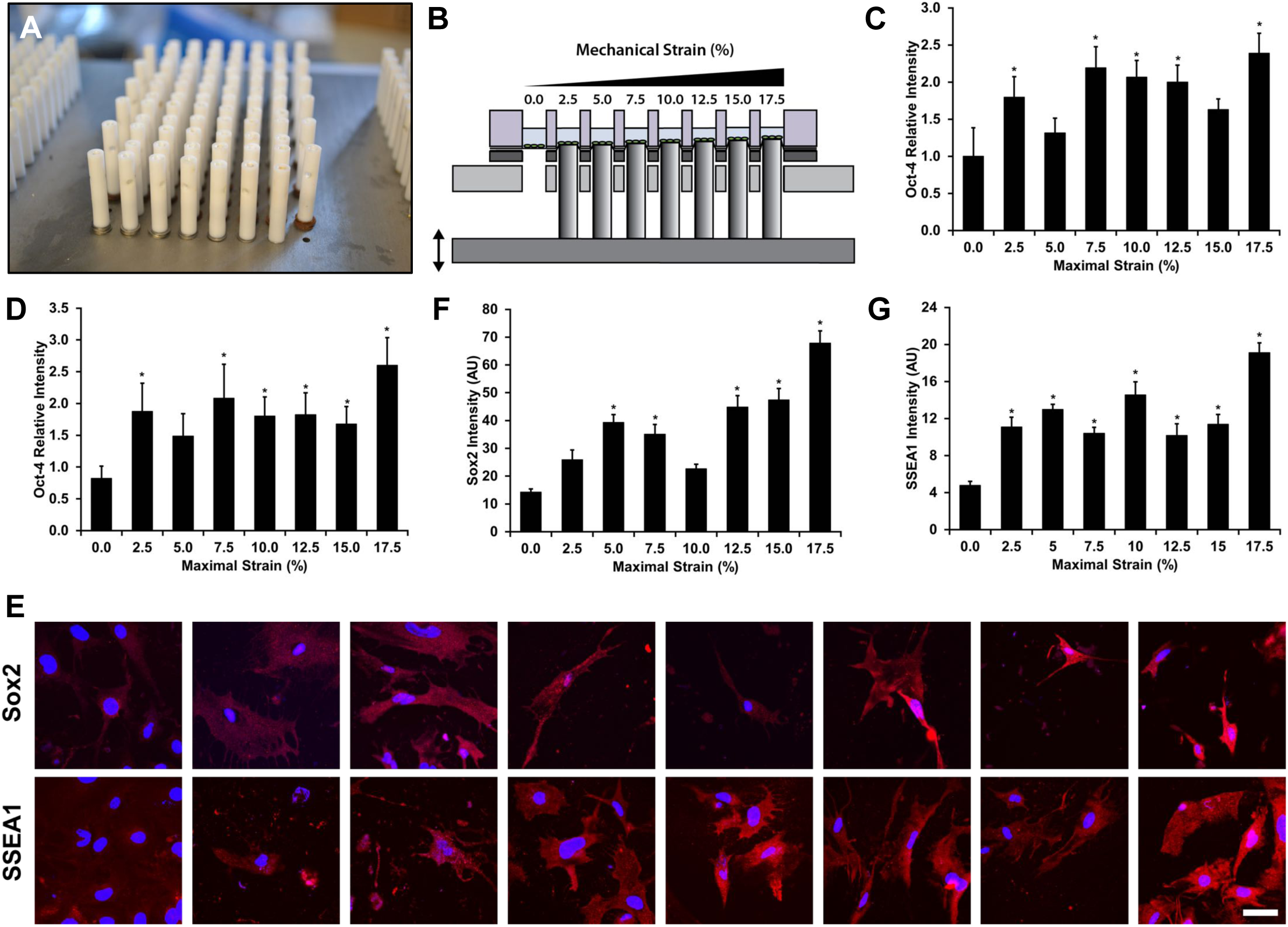
Mechanical strain enhances pluripotency transcription factors in mouse fibroblasts. (A) Thin shims were added to adjust the height of the pistons to apply varying mechanical strain across the plate. (B) Diagram of one row of pistons displacing the membranes. When the lowest piston is applying 2.5% strain the highest piston is applying 17.5% strain. Mechanical stain was applied at 0.1 Hz for four hours per day for seven days. (C) Expression of Oct-4 eGFP in MEFs after one day of mechanical loading at varying levels of strain. ^*^*p* < 0.05 versus static control group. (D) Oct-4 eGFP signaling in MEFs after seven days of mechanical load. ^*^*p* < 0.05 versus static control group. (E) Immunostaining for Sox2 and SSEA1 in MEFs treated with mechanical stain for 7 days. Bar = 50 µm. (F) Quantification of Sox2 in MEFs after 7 days of mechanical load. ^*^*p* < 0.05 versus static control group. (G) Quantification of SSEA1 in MEFs after 7 days of mechanical load. ^*^*p* < 0.05 versus static control group.

### High throughput mechanobiological screen of small molecule inhibitors combined with mechanical load to enhance Oct-4 expression in MEFs

Using the high throughput aspect of the loading system, we performed a screen in which we applied 17.5% strain to Oct-4 eGFP MEFs at 0.1 Hz for 14 days. We applied the mechanical load using either a sinusoidal waveform or a brachial waveform that has a shape that mimics the strain in the brachial artery during the cardiac cycle. In addition, we treated the cells with a library of small molecule kinase inhibitors. Using this screen, we could optimize the synergy between mechanical and pharmacological treatments to maximize Oct-4 expression in MEFs. Under pharmacological treatment in static conditions, many of the kinase inhibitors decreased Oct-4 expression below baseline levels (already very low) in the MEFs (**Fig. 3**). Cells treated with sine waveform and kinase inhibitors had predominantly either no alteration in Oct-4 or a reduction in Oct-4 expression. Under brachial waveform loading, there was increase in Oct-4 expression with most treatments. Notably, treatment with DMSO increased the Oct-4 expression in combination with brachial waveform mechanical loading in comparison to control cells with brachial loading (**Fig. 3**). In addition, several of the kinase inhibitors increased the Oct-4 expression significantly including a PKC? inhibitor (Enzastaurin; CAS 170364-57-5), a β-Catenin/Tcf Inhibitor (FH535; CAS 108409-83-2), and a GSK-3K inhibitor (SB-431542; CAS 280744-09-4), in combination with brachial loading.

**Figure 3.**
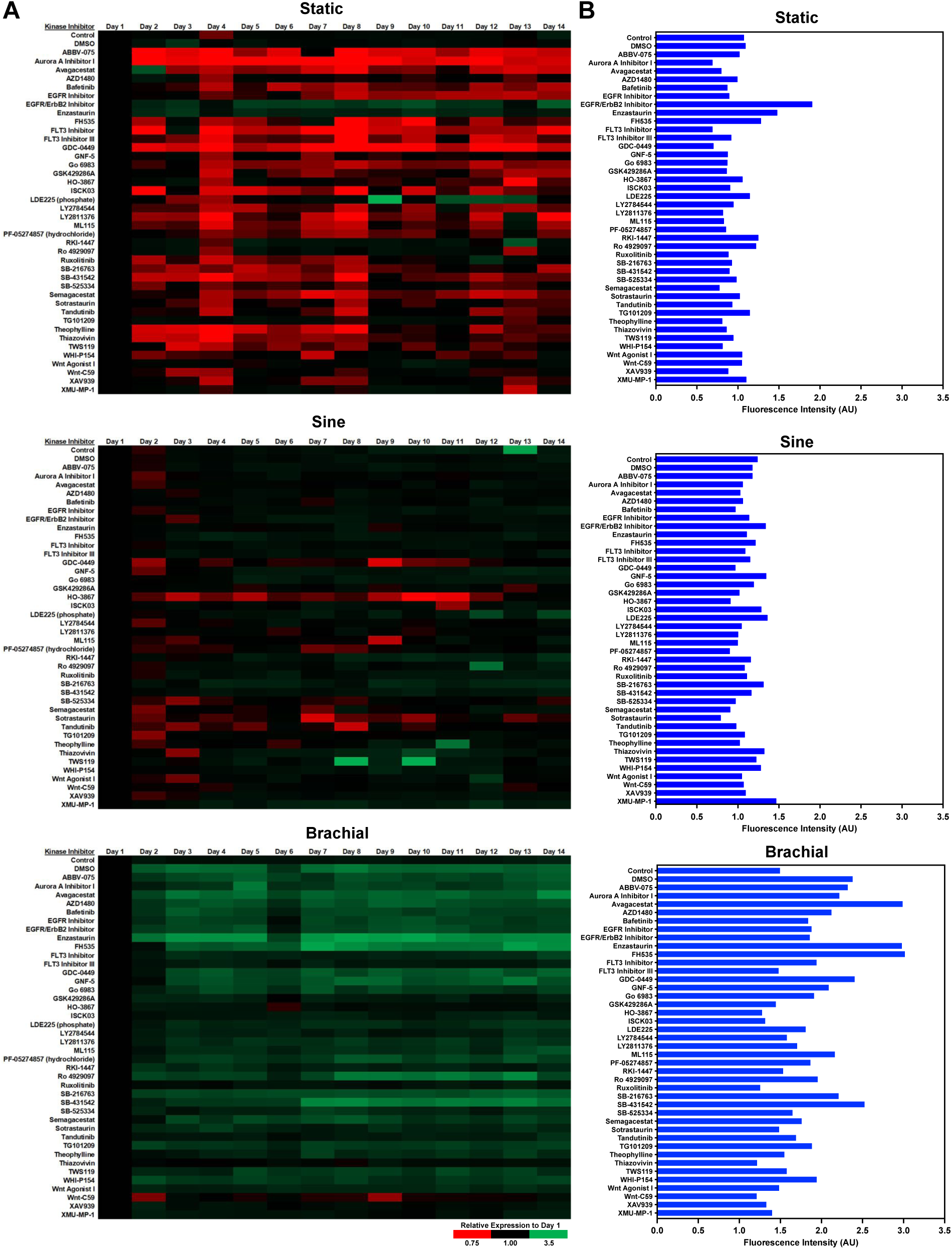
High throughput mechanobiological screen for small molecule inhibitors that synergistically increase Oct-4 GFP with mechanical loading. The MEFs were treated with 17.5% mechanical strain at 0.1 Hz for four hours per day for 14 days in the presence of compounds from a library of kinase inhibitors. The expression of Oct-4 GFP was measured using a plate reader each day. (A) Heat maps of Oct-4 GFP fluorescence for the various treatments. (B) Mean values of Oct-4 GFP expression after seven days of treatment.

### Optimized mechanical and pharmacological conditioning increases the expression of pluripotency related transcription factors and cellular marker in MEFs

To further confirm the expression of pluripotency related markers in the MEFs, we treated MEFs with a subset of conditions from our high throughput screen and then further analyzed these cells for the expression of pluripotency related transcription factors and gene expression for markers related to different stages of pluripotency (Schwarz et al., 2018). We found significant increases in SOX2 and NANOG with the brachial waveform and kinase inhibitors by immunostaining (**Fig. 4A, B**). In addition, we performed PCR for a broader set of markers relating to iPSC development from MEFs. Pluripotency-related transcription factors were increased by brachial loading and kinase inhibitor treatments including OCT-4, NANOG and SOX2 (**Fig. 4C**). We observed a reduction in mRNA expression for VCAM1 for the majority of the treatments but no significant change in PDGFRB (both MEF markers; **Fig. 4C**). We also observed no significant changes in the iPSC marker EPCAM among the groups (**Fig. 4C**).

**Figure 4.**
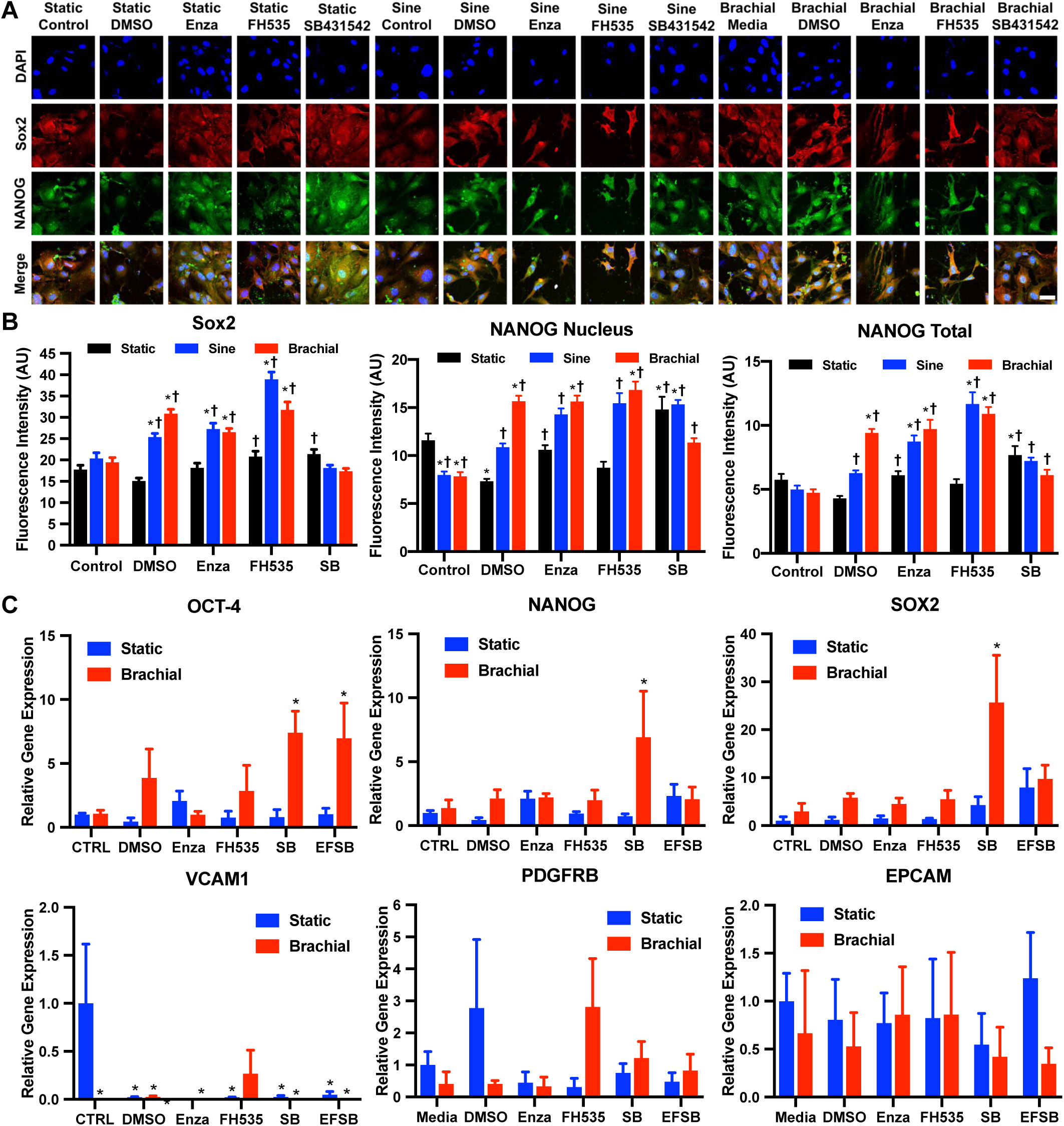
Optimized mechanical and pharmacological conditioning increases the expression of pluripotency markers in MEFs. (A) Immunostaining for Sox2 and NANOG in MEFs loaded under static, sinusoidal, or brachial waveform at 0.1 Hz, 17.5% maximal strain for 4 hours a day for 14 days. Bar = 100 µm. (B) Quantification of Sox2, NANOG nucleus, and NANOG total expression in MEFs after 14 days of mechanical load. ^*^*p* < 0.05 versus static control group. ^†^*p* < 0.05 versus static DMSO group. (C) Gene expression of pluripotency, MEF, and iPSC marker in MEF after 7 days of mechanical load measured through real-time PCR. ^*^*p* < 0.05 versus static control group.

## Discussion

While iPSCs have immense potential for their ability to mimic disease in vitro, there remain concerns for the use of genetically modified cells as therapeutics in patients. Thus, reprogramming strategies that do not use genetic modification would be highly advantageous for enhancing the therapeutic potential of iPSCs. Several studies have found that small molecule inhibitors can substitute for one or more pluripotency transcription factors (Hou et al., 2013; Ichida et al., 2009; Lin et al., 2009; Zhu et al., 2010). However, prior studies have found that mechanical forces can both inhibit and enhance the development of pluripotency. In this study, we developed a novel mechanical screening system that enabled a more complete exploration of mechanical conditions that could potentially alter the development of pluripotency in somatic cells. Using this screening system, we found optimal mechanical conditions that enhanced the expression of pluripotency transcriptions factors and that this could be further enhanced through treatment with small molecule inhibitors.

In our study, high levels of mechanical strain were needed to induce pluripotency transcription factor expression. In addition, the use of a physiologic waveform (brachial) with a high strain rate was most effective at inducing many of the transcription factors. The expression of pluripotency transcription factors in MEFs was further increased by adding in inhibitors to PKCβ, β-catenin/Tcf, or TGF-βRI/ALK4/ALK7 in combination with the optimal mechanical loading conditions. Mechanical strain alone was able to increase OCT-4 eGFP expression by nearly 2.5 fold. This effect was present after one day of loading and increased slightly over time for the seven days. With the addition of specific inhibitors with mechanical load, the enhancement of OCT-4 eGFP was increased to 3.2 fold over baseline levels. The inhibitor SB431542 in combination with brachial load enhanced OCT-4, SOX2, SSEA1 and c-Myc. In contrast, the inhibitor FH535 in combination with brachial loading was the best condition for increasing NANOG gene expression. While there was consistent increases in pluripotency transcription markers, gene expression for other markers of iPSC generation were mixed. We observed a reduction in some of the MEF markers (VCAM1) but observed no change in another marker (PDGFRB). In addition, we observed no change in the iPSC marker EPCAM. Thus, the effects of mechanical conditioning/small molecule treatment enhance the pluripotency transcription factors but do not completely alter the marker expression of the MEFs to match the development of iPSCs.

Our study also provides a major technical advancement through the development of a flexible, high throughput format for the study of the mechanical conditioning of stem cells. Systems for studying applied mechanical forces to cultured cells almost universally use flexible cell culture surfaces that can be expanded through applied mechanical forces (Brown, 2000; Davis et al., 2015; MacQueen et al., 2013). As the membrane is expanded, the cells are exposed to mechanical strain through their coupling to the flexible membrane. The expansion of the membrane can be induced through the displacement of a piston, (Baker et al., 2008; Schaffer et al., 1994) pneumatic suction (Zhong et al., 1998) or direct traction on the membrane (Shao et al., 2013). Systems using a rotational motor with a cam to drive piston motion are limited in that they can apply only one waveform without modifying the cam shape (albeit at varying frequencies) (Schaffer et al., 1994). Pneumatic systems are commercially available but also have limitations in the dynamics of the pneumatic system and ability to apply only a single strain at one time (Colombo et al., 2008). There have only been limited studies on making these systems high throughput and few systems are currently capable of applying more than a single magnitude of mechanical strain simultaneously (Kamotani et al., 2008; Simmons et al., 2011). The HT-BOSS system overcomes many limitations of previous systems with increased throughput and the use of the standard 96 well format, enabling it to integrate with plate reader assays and high content imaging systems. In addition, the linear motor allows the application of arbitrary waveforms with highly controlled dynamics. The independent adjustment of the pistons also provides a means to apply multiple strain levels simultaneously and to calibrate the system to accurately apply mechanical load in each well.

Overall, our results demonstrate the ability of combined mechanical/pharmacological conditioning to prime the pluripotency of somatic cells in the absence of genetic modification. The induction of the transcription factors was highly dependent on the magnitude of mechanical strain and the complex dynamics of the applied waveform. Moreover, the response of the cells to drugs was highly dependent on the mechanical conditions, with compounds causing opposing response in static versus loaded conditions. Thus, mechanobiological screening may provide complementary enhancement of strategies for chemically inducing pluripotency and aid in the development of safe methods for creating iPSCs for therapy in human patients.

## Acknowledgements

The authors gratefully acknowledge funding through the American Heart Association (17IRG33410888), the DOD CDMRP (W81XWH-16-1-0580; W81XWH-16-1-0582) and the National Institutes of Health (1R21EB023551-01; 1R21EB024147-01A1; 1R01HL141761-01) to ABB.

## Author Contributions

JL, MO, PM, YU, LS and MW conducted the experiments. JL and ABB designed experiments and wrote the paper.

## Materials and Methods

### Cell culture

Mouse embryonic fibroblasts (PrimCells LLC) or Oct-4 eGFP reporter expressing MEFs (EMD Millipore) at passages 5 were cultured in 4.5 g/L D-glucose DMEM medium supplemented with 10% fetal bovine serum, L-glutamine, and penicillin/streptomycin. All cells were cultured at 37°C and 5% CO_2_. For passaging, 70% confluent MEFs were first washed with PBS. Cell were agitated with 0.25% EDTA-trypsin at 37 °C for 1 minute. Cell suspension was then collected and centrifuged at 500 g for 5 minutes. The cells were then seeded to a new substrate at 2,000 cells per cm^2^.

### Mechanical loading device

Strain was applied to cells using a custom made device that displaces pistons through the flexible culture surface. The device operates using a true linear motor that drives a platen on motion rails. Linear ball bearings are used along the rails to minimize friction, while fixed springs on the rods help reduce the load on the motor and prevent the platen from moving while the device is turned off. There are 576 individual PTFE pistons mounted on the platen that are removable and can be calibrated individually through thin shims. A top plate on the system has mounting holes for custom designed culture plates that hold a silicone membrane sandwiched between steel plates and silicone gaskets. The silicone membrane can be coated to allow culture of cells and the entire geometry matches that of a standard 96 well culture plate. Mechanical strain is applied when the pistons are moved into the silicone membrane, causing displacement and application of stretch to the cells. The linear motor is hygienically sealed and feedback controlled by software that regulates the current through the coils around the motor (Copley Motion). To avoid excess heat from the current generated and from high temperature inside cell incubators, a cooling system is integrated with an external water bath circulating chilled water into the motor’s enclosure.

### Flexible-bottom culture plate assembly

The cell culture plate was assembled by sandwiching custom made parts with the flexible silicone rubber to provide a cell culture substrate. The culture plate consists of a polycarbonate top plate and aluminum bottom plate that are held together by eight screws. Sandwiched between the top and bottom plate is a 0.005’’ thick silicone sheet (Specialty Manufacturing, Inc.) with a rubber gasket to prevent leaking. The plates are sterilized prior to cell seeding by UV light and are coated by treatment with 50 µg/mL fibronectin at 37°C overnight. On the day of mechanical loading, the plates were mounted to the top plate of the device with eight additional screws.

### Calibration of mechanical strain

Mechanical strain applied to the flexible membrane was measured by recording changes in the marks drawn on the membrane. A uniform array of marks was created on the cell culture surface using a stencil and silicone glue. The pistons were displaced at small increments through the membrane and a high magnification image of the mark was recorded using a high-resolution camera (Basler AG). The displacement of the membrane was measured using Metamorph Imaging software. For dynamic mechanical loading, strain was measured by recording a video at 60 frames per second. The membrane was displaced in sinusoidal, aortic, and brachial waveforms created from clinical arterial distension data (Lee et al., 2013). For each waveform, three cycles of the waveform were recorded and averaged.

### Computational modeling of the system

Fluid mechanics in the cell culture media was modelled using finite element software (COMSOL). Briefly, a cylindrical shape fluid structure was used to model a single well with viscosity and density of standard DMEM media. The bottom surface was displaced in a sinusoidal motion over time with three frequencies (0.1Hz, 1Hz, and 2Hz) in combination with three maximum magnitudes (1%, 5%, and 10%). A series of mesh and tolerance optimization were performed to optimize these parameters. The maximum and average fluid velocities at various cross sections were computed. Average shear stress in various locations and over various time points were computed as well with the focus on the location of the bottom surface undergoing displacement where the cells are located.

### Immunostaining

Following the treatments, the cells were fixed in 4% paraformaldehyde in PBS for 10 minutes followed by washing and permeabilization with 0.1% Triton X-100 PBS for 5 minutes. Next, samples were blocked with PBS containing 5% FBS and 1% BSA for 40 minutes. After washing, cells were incubated with primary antibodies at 1:100 dilution ratio in PBS with 1% BSA overnight at 4°C. Primary antibodies used include NANOG (ab80892; Abcam), Sox2 (4744S; Cell Signaling Technology), and SSEA-1 (ab79351; Abcam). The samples were then washed twice in PBS with 1% BSA and incubated with secondary antibodies at 1:1000 dilution ratio in PBS with 1% BSA for 2 hours in a light protected environment. Cells were then washed with PBS with 1% BSA prior to mounting in anti-fade media (Vector Laboratories, Inc.). The samples were imaged using FV10i Confocal Laser Scanning Microscope (Olympus, Inc.). Images of the fluorescent cell cultures were then traced in Adobe Photoshop for fluorescence intensity quantification.

### Real Time PCR

For gene expression analysis, mRNA was purified and collected from the cells using RNEasy Mini Kit (Qiagen, Inc.). Reverse transcription was performed using TaqMan (Thermo Fisher) or QuantiTect (Qiagen) reverse transcription reagents. PCR was performed using SYBR Green PCR Master Mix (Applied Biosystems) with ViiA7 Real-Time PCR System (Thermo Fisher). Custom primers used for real time PCR are listed in **Supplemental Table 1**. For analysis, 18s ribosomal RNA (18s rRNA) was used as a house keeping gene.

### Multi-strain mechanical loading and measurement of Oct-4 GFP expression

To determine the optimal strain for Oct-4 expression in MEF, Oct-4 eGFP reporter expressing MEFs were exposed to either static or sinusoidal waveform loading for 8 hours a day for 7 days using the multi-strain configuration of HT-BOSS, where the cells were exposed to maximal strain ranging from 0.0% to 17.5%, in 2.5% intervals. For measuring the expression of Oct-4 eGFP, plate reader was used (Varioskan; Thermo Fisher). Briefly, culture media was replaced with Tyrode’s Solution every day prior to the fluorescence measurement. After plate reading, the Tyrode’s solution was replaced with cell culture media.

### Kinase Inhibitor Drug Library

For the kinase screening study, Oct-4 eGFP expressing MEFs were cultured under static, sinusoidal waveform, or brachial waveform loading for 4 hours per day for 14 days at 0.1Hz and 17.5% maximal strain. During loading, the cells were treated with media no treatment, 1 µM DMSO, or one of the 40 kinase inhibitors listed in **Supplemental Table 2** at 1 µM (Cayman Chemicals). Each day, the cell culture media were replaced with Tyrode’s Solution and the Oct-4 GFP expression was measured using a plate reader. The Tyrode’s solution was replaced with media containing treatment every day prior to stretching.

### Statistical analysis

All results are shown as mean ± standard error of the mean. Multiple comparisons between groups were analyzed by two-way ANOVA followed by a Tukey post-hoc or a Dunnett post-hoc test when testing multiple comparisons versus a control group. A *p*-value of 0.05 or less was considered statistically significant.

## Supplementary Figure Legends

**Supplemental Figure 1.**
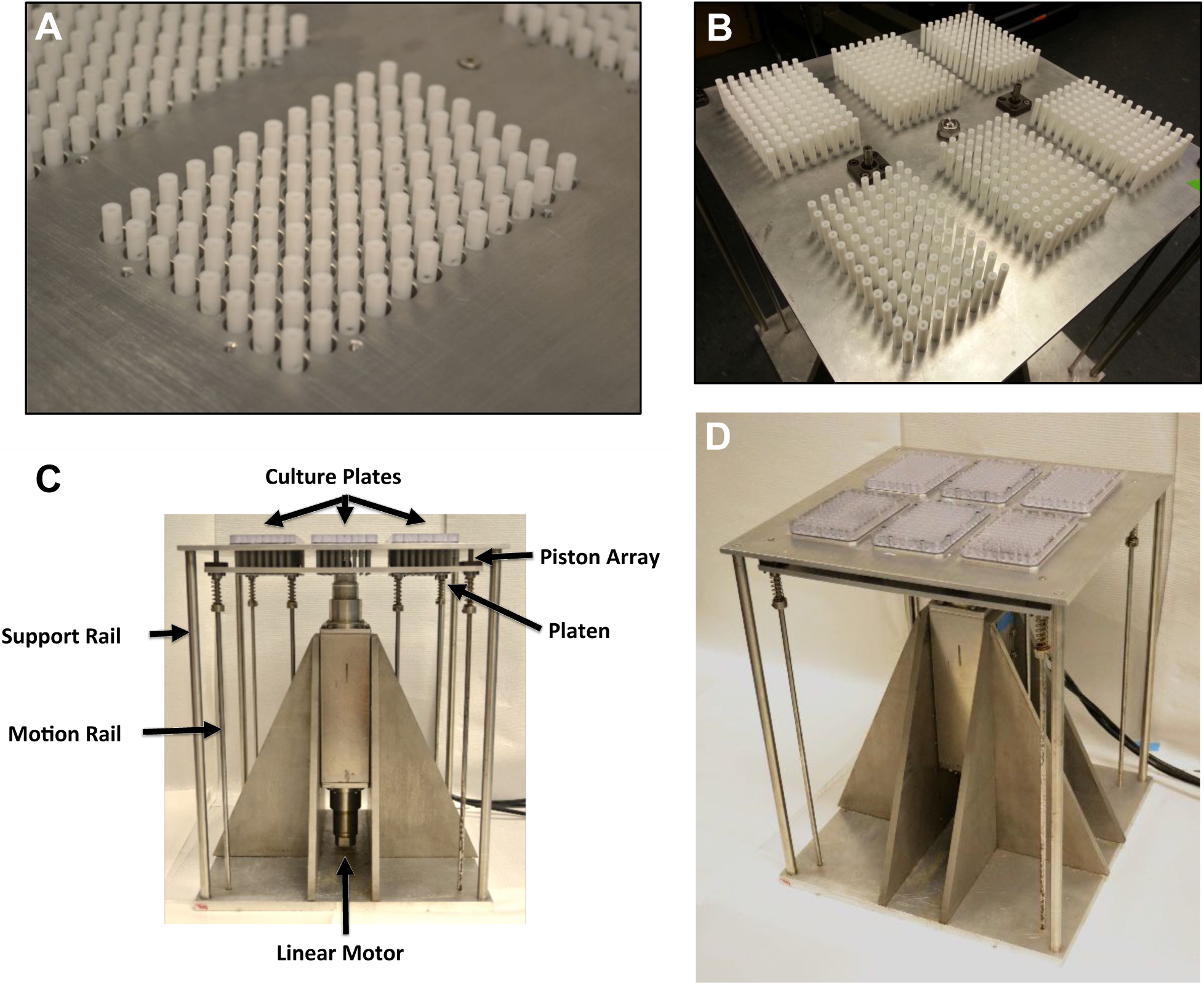
Assemble HT-BOSS system for applying mechanical strain. (A) A photograph of the PTFE pistons passing throughput a steel top plate. (B) The platen holds 576 pistons to apply load to six 96-well plates. (C) Overview of the system with culture plates mounted. (D) Picture of the completed system.

**Supplemental Figure 2.**
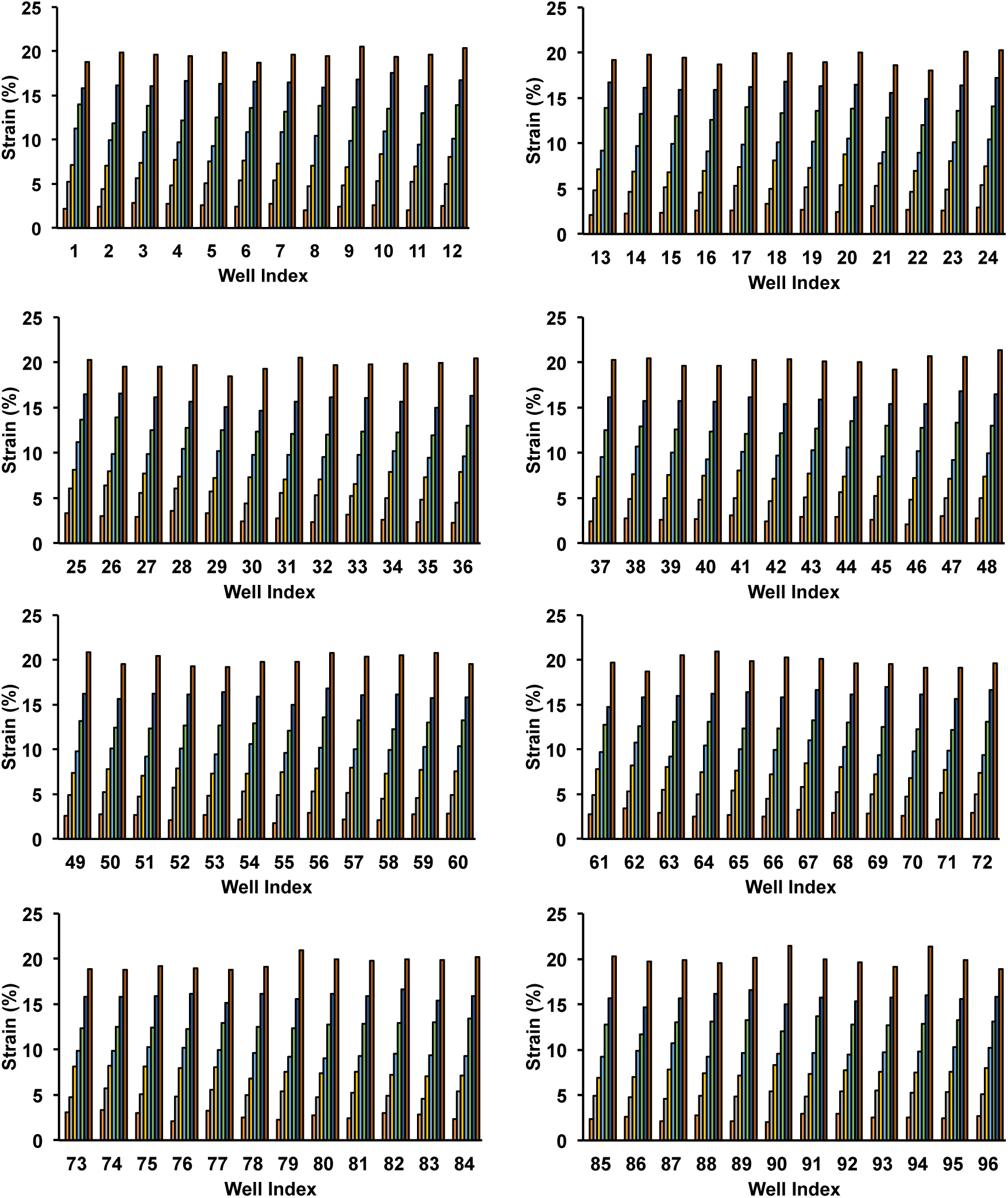
Detailed calibration of device to apply consistent mechanical load between culture wells. Well-by-well measurement of strain for a 96 well plate under load application.

**Supplemental Figure 3.**
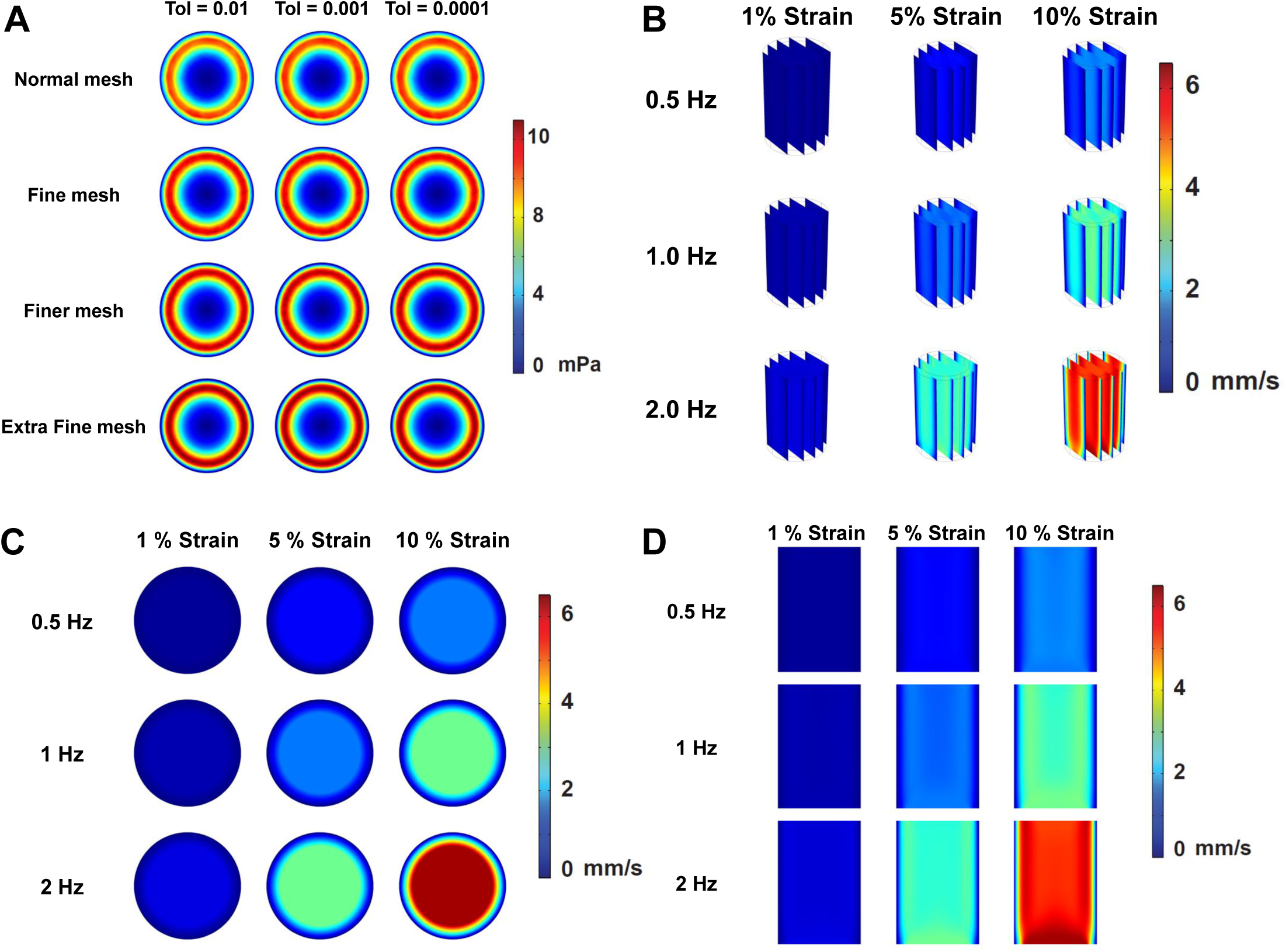
Optimization of computational mesh and peak flow rates in the computational model. (A) Optimization of the mesh size and tolerance for the simulation. (B-D) Peak flow rates within the well within the well during the application of mechanical strain.

**Supplemental Figure 4.**
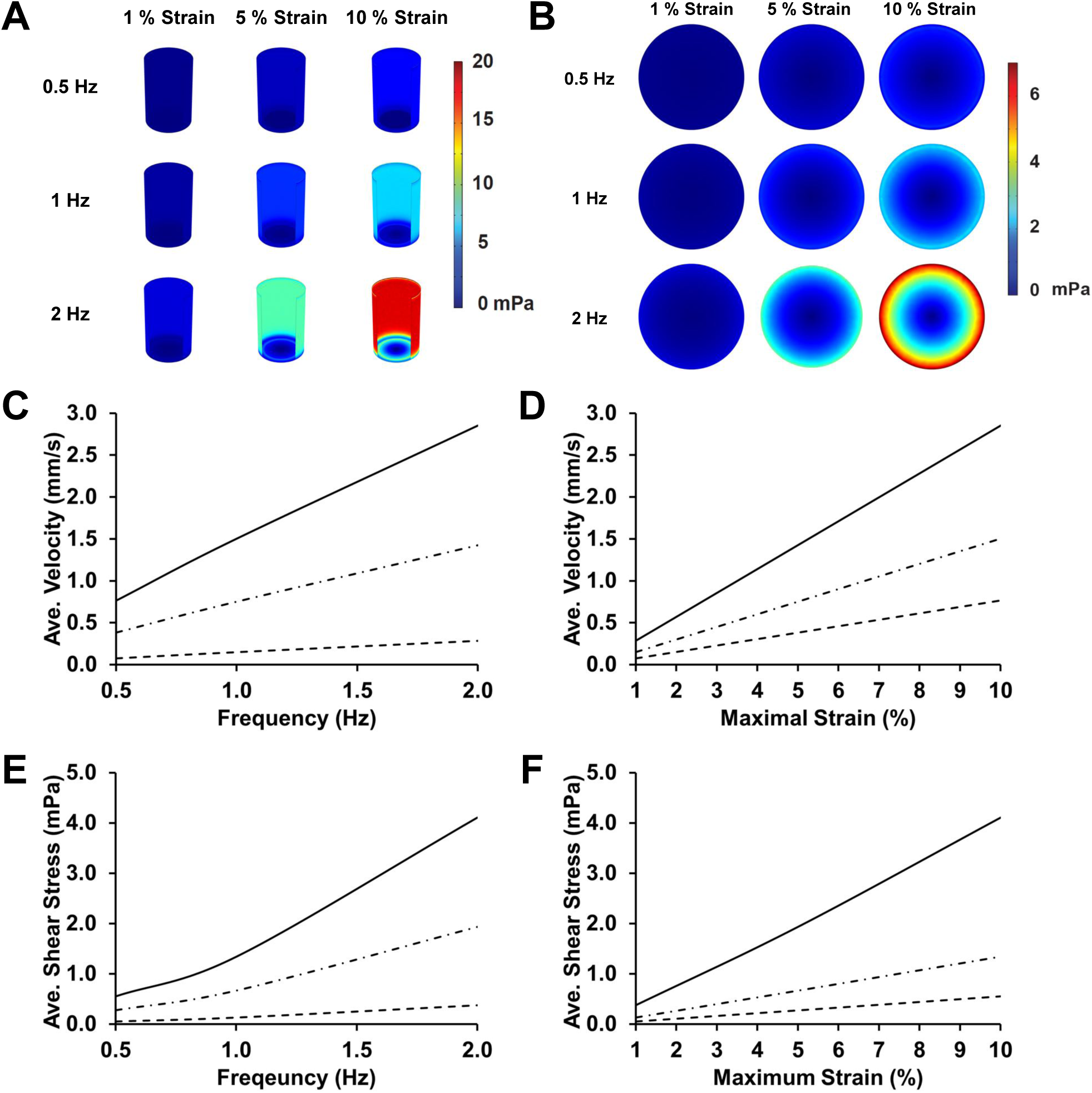
Computational modeling of fluid flow during mechanical loading. (A) Peak shear stress within the well during the displacement of the membrane. (B) Peak shear stress on the culture surface during mechanical loading. (C) Average fluid velocity within the well as a function of the frequency of loading. (D) Average fluid velocity within the well as a function of the maximal strain of loading. (E) Average shear stress on the culture surface as a function of the frequency of loading. (F) Average shear stress on the culture surface as a function of the maximal strain.

**Supplemental Figure 5.**
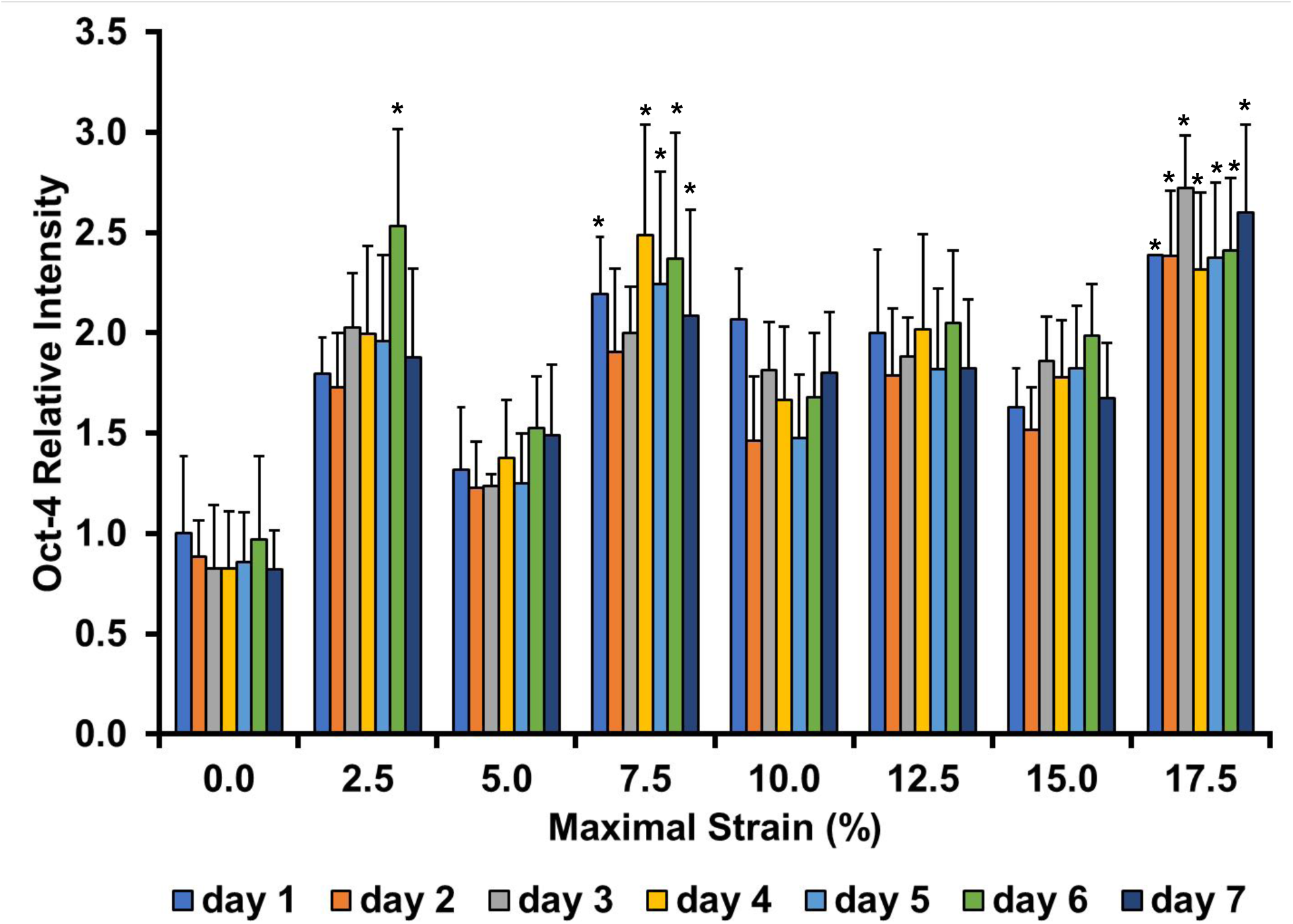
Time course of Oct-4 expression in MEFs treated with mechanical loading. Expression of Oct-4 eGFP in MEFs treated with mechanical strain at 0.1 Hz for four hours per day. ^*^*p* < 0.05 versus static day 1 group.

## Supplemental Tables

**Table 1.**
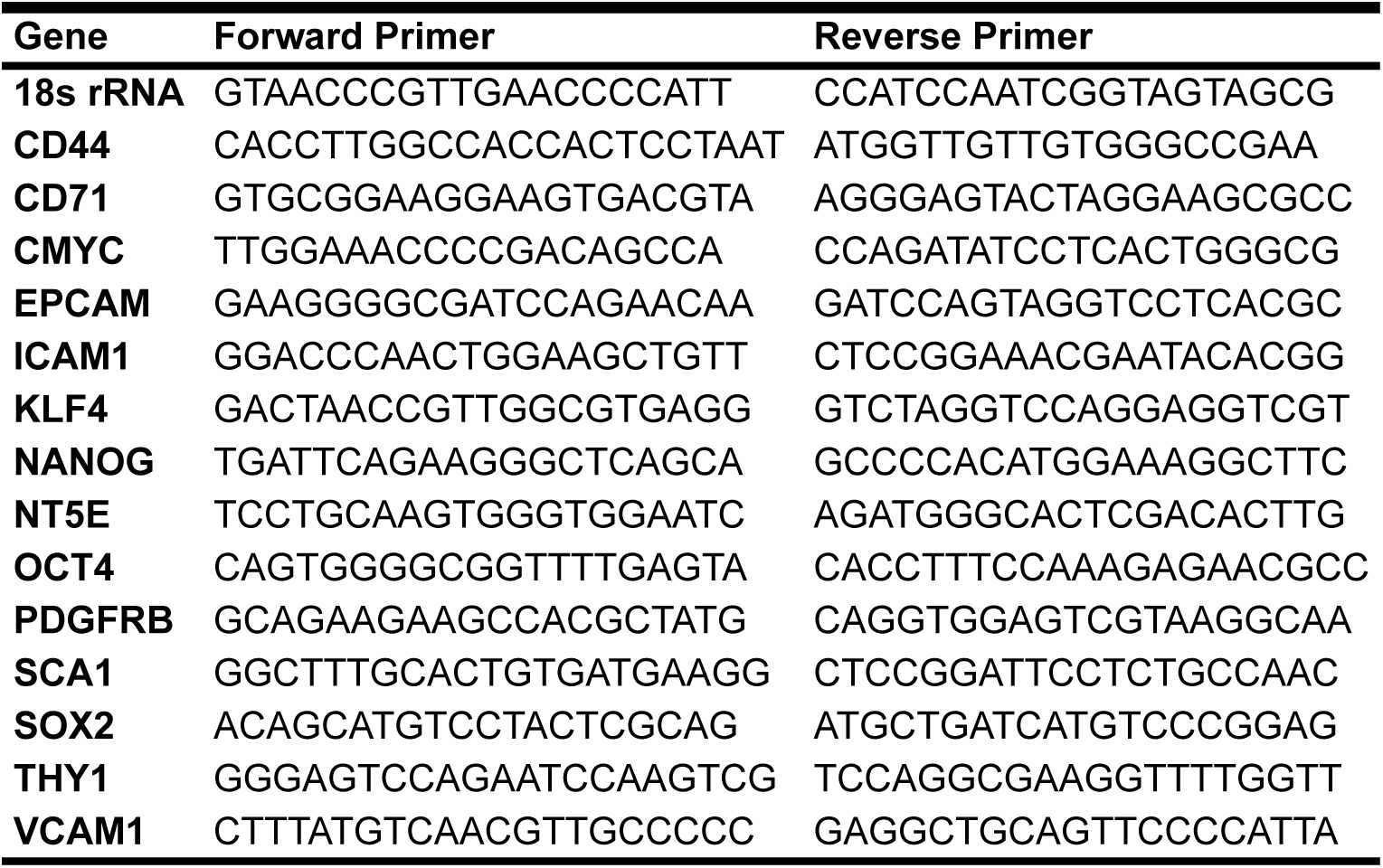
Primers used for PCR Analysis

**Supplemental Table 2.**
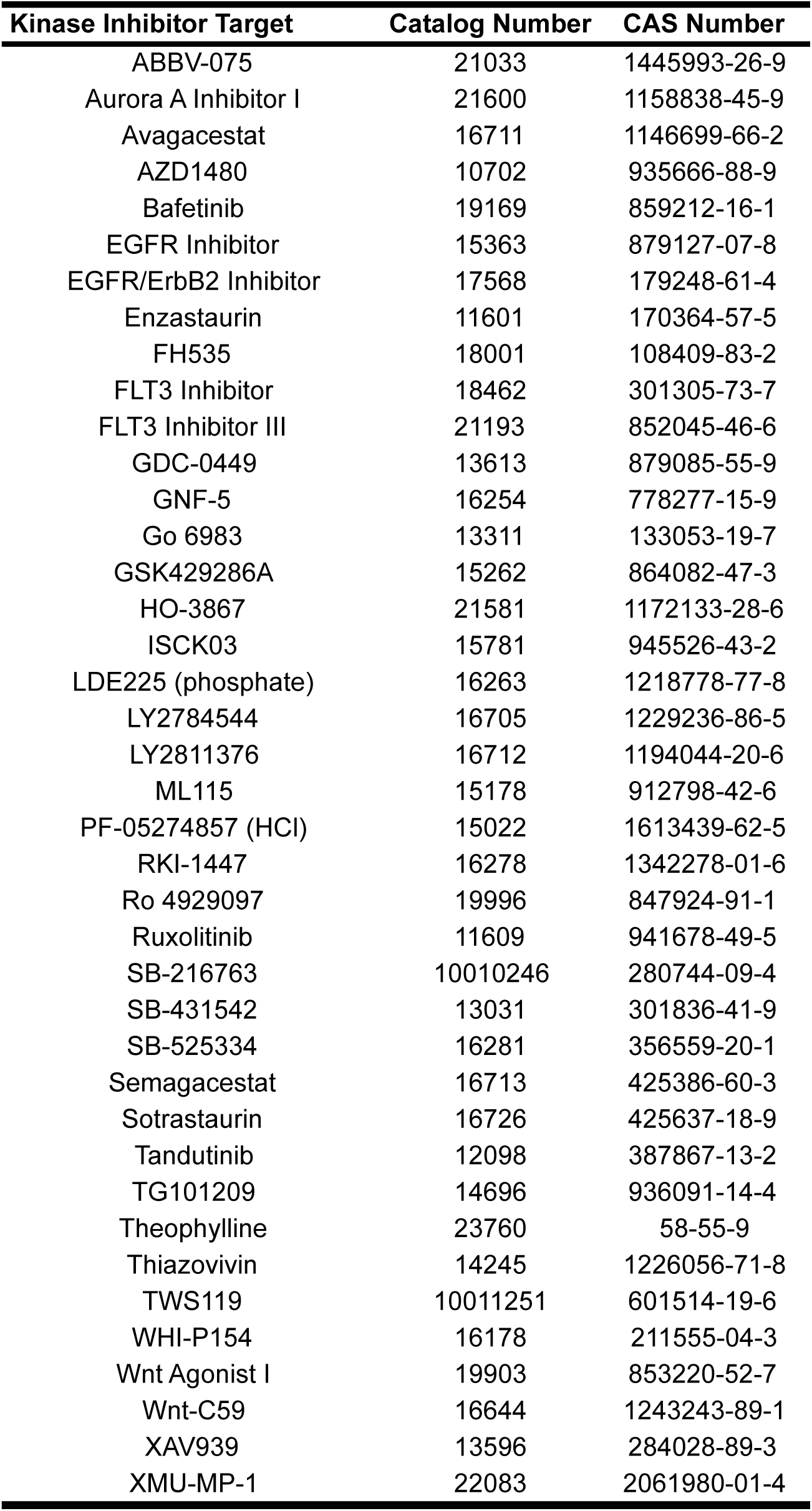
Kinase Inhibitors Used in the Studies

| **Kinase Inhibitor Target** | **Catalog Number** | **CAS Number** |
| --- | --- | --- |
| ABBV-075 | 21033 | 1445993-26-9 |
| Aurora A Inhibitor I | 21600 | 1158838-45-9 |
| Avagacestat | 16711 | 1146699-66-2 |
| AZD1480 | 10702 | 935666-88-9 |
| Bafetinib | 19169 | 859212-16-1 |
| EGFR Inhibitor | 15363 | 879127-07-8 |
| EGFR/ErbB2 Inhibitor | 17568 | 179248-61-4 |
| Enzastaurin | 11601 | 170364-57-5 |
| FH535 | 18001 | 108409-83-2 |
| FLT3 Inhibitor | 18462 | 301305-73-7 |
| FLT3 Inhibitor III | 21193 | 852045-46-6 |
| GDC-0449 | 13613 | 879085-55-9 |
| GNF-5 | 16254 | 778277-15-9 |
| Go 6983 | 13311 | 133053-19-7 |
| GSK429286A | 15262 | 864082-47-3 |
| HO-3867 | 21581 | 1172133-28-6 |
| ISCK03 | 15781 | 945526-43-2 |
| LDE225 (phosphate) | 16263 | 1218778-77-8 |
| LY2784544 | 16705 | 1229236-86-5 |
| LY2811376 | 16712 | 1194044-20-6 |
| ML115 | 15178 | 912798-42-6 |
| PF-05274857 (HCl) | 15022 | 1613439-62-5 |
| RKI-1447 | 16278 | 1342278-01-6 |
| Ro 4929097 | 19996 | 847924-91-1 |
| Ruxolitinib | 11609 | 941678-49-5 |
| SB-216763 | 10010246 | 280744-09-4 |
| SB-431542 | 13031 | 301836-41-9 |
| SB-525334 | 16281 | 356559-20-1 |
| Semagacestat | 16713 | 425386-60-3 |
| Sotrastaurin | 16726 | 425637-18-9 |
| Tandutinib | 12098 | 387867-13-2 |
| TG101209 | 14696 | 936091-14-4 |
| Theophylline | 23760 | 58-55-9 |
| Thiazovivin | 14245 | 1226056-71-8 |
| TWS119 | 10011251 | 601514-19-6 |
| WHI-P154 | 16178 | 211555-04-3 |
| Wnt Agonist I | 19903 | 853220-52-7 |
| Wnt-C59 | 16644 | 1243243-89-1 |
| XAV939 | 13596 | 284028-89-3 |
| XMU-MP-1 | 22083 | 2061980-01-4 |

